# Effect of diets containing whole or ground corn with and without soybean hulls on feeding calves during the suckling phase

**DOI:** 10.1101/486258

**Authors:** Aline Evangelista Machado Santana, José Neuman Miranda Neiva, Vera Lúcia de Araújo Bozorg, João Restle, Ubirajara Bilego

**Author notes:** Corresponding author (AEMS). These authors contributed equally to this work (Aline Evangelista Machado Santana, José Neuman Miranda Neiva). These authors also contributed equally to this work (Vera Lúcia de Araújo Bozorg, João Restle, Ubirajara Bilego).

## Abstract

This work evaluated the use of soybean hulls and whole or ground corn in the diets of suckling calves. Diets containing two levels of soybean hull inclusion (0 and 400.1 g/kg) and corn in different physical forms (whole or ground) were evaluated in the diets of newborn crossbred dairy calves that were housed and received experimental diets plus four liters of milk per day over 56 days. Weekly samples of food, diets and leftovers were taken to determine dry matter and nutrient intake. To evaluate the apparent digestibility, samples were taken from the diets, leftovers and feces for three consecutive days using titanium dioxide as an indicator. Blood samples were also collected to evaluate the blood indicators. Including soybean hulls in the diet increased the consumption of neutral detergent fiber but reduced the consumption of non-fibrous carbohydrates, which was also reduced by using whole corn in the diet. Total digestible nutrient consumption did not vary, although its value was reduced by using whole corn and including soybean hulls. The apparent digestibilities of the dry matter and crude protein were similar, resulting in similar performances between the animals, regardless of the factors analyzed. Using soybean hulls or whole corn did not affect blood indicators or feeding costs. Soybean hull or whole corn usage did not affect the performance of crossbred dairy calves during rearing.

## Introduction

Although milk producers are efficient at producing heifers, male cattle production is typically neglected. In Brazil, the number of cows milked corresponds to 23 million animals [1] with a birth rate of 50% male; thus, 11.5 million calves are produced annually. Although no official data exist on the number of calves slaughtered shortly after birth in Brazil, approximately 40% of these calves are slaughtered in their first days of life, representing approximately 4.6 million calves. This occurs because producers consider that these calves will increase their production costs without favorable financial returns; however, these animals can yield good productive results when their nutritional and hygiene requirements are fulfilled [2], mainly during the first days of the animal’s life (0 to 60 days).

This period is the most costly in the production system; thus, solutions are needed to balance these costs, such as using palatable solid foods, which stimulate the animals to eat. This allows the animals’ weaning age to be reduced because they benefit from more efficient use of the nutrients in solid food [3,4], which reduces the problems caused by weaning the animals early [5-7], increases the amount of milk available for sale and reduces the demand for labor.

However, the use of feedstuff, such as corn and soybeans meal, which are traditionally used in animal feed, in the biofuel production, has increased the price of these inputs [8]. Thus, alternatives to these foods are needed to reduce feed costs, such as by-products including soybean hulls (SH), which are a good quality food with a low market value [9]. In addition, the degree of feed processing can be reduced by supplying whole corn grain, which reduces processing costs and allows supplying solid food with large particle sizes to induce the animals’ normal chewing behaviors [10].

This study haved as objective evaluated the effect of diets containing soybean hulls and whole or ground corn on the digestibility, cost of diets and performance of crossbred dairy calves during the rearing phase.

## Materials and methods

The experimental procedures were approved by the Animal Ethics Committee of the Federal University of Tocantins under case no. 23101.004142/2015-06, which considers the ethical guidelines set by the Council of International Organizations of Medical Sciences, CIOMS/OMS, 1985, for research involving animals. The experiment was performed in the municipality of Rio Verde, Goiás state, Brazil, located at 17°47’52” south latitude and 50°55’40” west longitude, from June to August 2013. The local climate is “Aw”, per the Köppen-Geiger classification, and the precipitation and mean temperature in the months of the experiment were 0.23 mm and 22.08°C, respectively. The relative humidity was 61.42%.

The experiment was conducted in a completely randomized design with the treatments distributed in a 2 × 2 factorial arrangement without soybean hull (NSH) or with 400.1 g/kg soy hull (WSH) and corn in two forms, whole (WC) or ground (GC), with seven replicates. Twenty-eight newborn crossbred calves were used with bloodlines varying from 3/4 to 5/8 Holstein × Zebu and initial weights of 33.01 kg. The total experimental period was 60 days; the calves were fed colostrum for 4 days, and data collection occurred on the remaining 56 days.

The animals were treated against endo and ectoparasites, received injectable vitamins A, D and E and were housed in individual shelters covered with a shaded area of 2 m^2^ and individual feeders and drinking troughs with rice husk bedding. The bedding was stirred daily by removing the wet bedding and feces, and each bed was replaced every seven days.

During the experimental period, the animals received diets composed of milk and the concentrate feed, which was formulated as isoprotein and contained corn, soybean hull, soybean meal and a commercial mineral core (Table 1).

**Table 1.** Chemical and bromatological food compositions used to formulate the concentrates

| Variables, g/kg DM | Corn | Soybean Meal | Soybean Hull |
| --- | --- | --- | --- |
| Dry Matter<sup>1</sup> | 850.1 | 830.3 | 889.7 |
| Ashes | 17.1 | 65.1 | 45.8 |
| Neutral Detergent Fiber | 113.0 | 154.5 | 655.6 |
| Hemicellulose | 83.5 | 41.8 | 261.9 |
| Fiber in Acid Detergent | 29.5 | 112.7 | 393.7 |
| Cellulose | 18.8 | 100.0 | 359.8 |
| Lignin | 10.7 | 12.7 | 33.9 |
| Ether Extract | 41.5 | 21.3 | 10.5 |
| Crude Protein | 81.9 | 474.9 | 146.9 |
| NDIN<sup>2</sup> | 83.8 | 7.0 | 381.4 |
| ADIN<sup>3</sup> | 47.9 | 3.8 | 139.0 |
| Non-fibrous Carbohydrates | 746.5 | 284.2 | 141.2 |
| Total Carbohydrates | 859.5 | 438.7 | 796.8 |

<sup>1</sup>Dry matter: in g/kg of natural matter; <sup>2</sup>NDIN: neutral detergent insoluble nitrogen in g/kg in total nitrogen;
<sup>3</sup>ADIN: acid detergent insoluble nitrogen in g/kg of total nitrogen.

The concentrate (Table 2) was supplied to the animals ad libitum, keeping 5% of the leftovers, which were removed once weekly. Samples were then collected from the leftovers, while food and concentrate samples were collected during their preparation. Milk was supplied in a restricted manner so that the animals consumed 4 liters per day, divided into two feeding times, with 31.8 g/kg ether extract (EE), 29.5 g/kg crude protein (CP), 45.2 g/kg lactose, 110.7 g/kg total dry extract and 79.8 g/kg fat-free dry extract.

**Table 2.** Ingredient proportions and chemical-bromatological composition of the concentrates

| Food, g/kg DM | Ground corn |  | Whole Corn |  |
| --- | --- | --- | --- | --- |
|  | NSH<sup>1</sup> | WSH<sup>2</sup> | NSH | WSH |
|  | Proportion of Ingredients |  |  |  |
| Whole Corn | - | - | 712.0 | 400.1 |
| Ground Corn | 712.0 | 400.1 | - | - |
| Soybean Hull | - | 400.1 | - | 400.1 |
| Soybean Meal | 238.0 | 148.0 | 238.0 | 148.0 |
| Mineral Core<sup>3</sup> | 50.0 | 50.0 | 50.0 | 50.0 |
| Variables, g/kg DM | Chemical-bromatological Composition |  |  |  |
| Dry Matter<sup>4</sup> | 890.6 | 890.1 | 897.8 | 909.0 |
| Ashes | 71.9 | 82.7 | 87.5 | 99.3 |
| Neutral Detergent Fiber | 123.0 | 381.5 | 110.2 | 379.8 |
| Hemicellulose | 81.3 | 188.1 | 70.8 | 191.9 |
| Acid Detergent Fiber | 41.7 | 193.4 | 39.4 | 187.9 |
| Cellulose | 30.9 | 173.6 | 28.5 | 168.1 |
| Lignin | 10.8 | 19.8 | 10.9 | 19.8 |
| Ether Extract | 33.7 | 27.6 | 34.2 | 28.4 |
| Crude Protein | 176.5 | 176.8 | 175.9 | 173.5 |
| NDIN<sup>5</sup> | 89.6 | 87.9 | 84.8 | 86.8 |
| ADIN<sup>6</sup> | 43.8 | 42.9 | 42.4 | 44.6 |
| Non-fibrous Carbohydrates | 594.9 | 331.4 | 592.2 | 319.0 |
| Total Carbohydrates | 717.9 | 712.9 | 702.4 | 698.8 |

<sup>1</sup>NSH: no soybean hull; <sup>2</sup>WSH: with soybean hull; <sup>3</sup>Mineral Core values: minimum calcium 220 g/kg, phosphorus 80 g/kg, vitamin A 300,000 IU, vitamin E 300 mg/kg, copper 860 mg/kg, zinc 3 g/kg, iodine 120 mg/kg, iron 2 g/kg, manganese 1350 mg/kg, selenium 20 mg/kg, cobalt 80 mg/kg and monensin 1200 mg/kg; <sup>4</sup>Dry matter: in g/kg of natural matter; <sup>5</sup>NDIN: neutral detergent insoluble nitrogen in g/kg of total nitrogen; <sup>6</sup>ADIN: acid detergent insoluble nitrogen in g/kg of total nitrogen.

Feed concentrate samples were predried in a forced ventilation oven at 65°C to determine dry matter (DM) (Method 934.01 [11]), crude protein (Method 979.09 [11]) and ash (Method 942.05 [11]). The levels of neutral detergent fiber (NDF), acid detergent fiber (ADF) and lignin were determined as described previously [12], and the percentage of ether extract (EE) was determined by washing with petroleum ether at 90°C for one hour [13]. The values of non-fibrous carbohydrates (NFC), total carbohydrates (TC) and total digestible nutrients (TDN) were calculated per [14], where TC = 100 - (%CP + %EE + % AS), NFC = TC – %NDF and observed TDN = CPD + (EED × 2.25) + TCD, where CPD = crude protein apparent digestible, EED = ether extract apparent digestible, AS = ashes and TDC = total carbohydrates digestible.

Dry matter intake (DMI), crude protein intake (CPI), ether extract intake (EEI), neutral detergent fiber intake (NDFI), non-fibrous carbohydrate intake (NFCI) and total digestible nutrient intake (TDNC) expressed in kilograms per day (kg/day) were calculated from the analyses. Average daily gain (ADG) and dry matter conversion efficiencies (DME) were calculated in kg of ADG /kg DM intake, crude protein efficiency (CPE) in kg ADG /kg of CP intake and total digestible nutrient efficiency (TDNE) in kg of ADG /kg of TDN intake. Total dry matter intake (TDMI), total crude protein intake (TCPI), total crude protein efficiency (TCPE) and total dry matter conversion efficiency (TDME) were calculated, which were considered the intake of DM and CP from milk intake.

At the beginning and end of the experimental period, the animals were weighed individually in the morning, without previous fasting, and these weights were considered the initial live weight (IW) and final live weight (FW). During these weighings, the following morphometric measurements were recorded: withers height (WH), rump height (RH), thoracic perimeter (TP), arm perimeter (AP) and body length (BL). These measurements were always performed on the animals’ left sides with the calves in quadrilateral position, using a hypometric measuring cane and a measuring tape in centimeters.

When the animals reached 56 days of age, food, leftover and fecal samples were collected and used to determine the apparent digestibility of the diets. These samples were collected after delivering 10 g of titanium dioxide daily (external indicator), supplied to the animals diluted in 50 ml of milk via esophageal catheter over 10 days, and stool was collected directly from the rectum during the last four days. These samples were immediately frozen at −20°C and subsequently transferred to the Laboratory of Animal Nutrition of the Federal University of Tocantins, where they were homogenized, thus forming composite samples that were predried in a forced ventilation oven at 65°C and analyzed to determine the total fecal production per the method described in [15] and used to calculate the apparent digestibility (DA) of the dry matter and the nutrients present in the diet, in which DA = 1 - [(ingested nutrient - excreted nutrient)/ingested nutrient.

At the end of the experimental period, blood samples were collected by venipuncture of the jugular vein, without prior fasting. To measure blood glucose, samples were collected in tubes containing 10 μl of EDTA and 10 μl of sodium fluoride. The samples were cooled and sent to the Animal Pathology Laboratory of the Cooperativa Agroindustrial dos Produtores Rurais do Sudoeste Goiano (Agroindustrial Rural Producers Cooperative of Southwest Goias; COMIGO), in Rio Verde-GO, where they were centrifuged for 15 minutes at 2000 g/min to separate the plasma and serum. These aliquots were placed in Eppendorf(®) tubes, identified and frozen at −20°C for subsequent analyses of glucose (GL), triglycerides (TG), total cholesterol (TCL), total protein (TP), albumin (ALB), urea (UR), aspartate aminotransferase (AST), alkaline phosphatase (ALT), and creatinine (CRT), which were performed at 37°C at the Laboratory of Animal Nutrition of the Federal University of Tocantins, using commercial tests from Labtest Diagnostica S.A.(®) and the Bioplus(®) Bio-2000 IL-A spectrophotometer.

To evaluate the costs associated with using the different diets, information was collected on the amount paid per kilogram of each food used to prepare the concentrates during the experimental evaluation period, during which time the value of the dollar corresponded to BRL 2.24. This information was used to calculate the cost per kilogram of diet (CKD), cost of daily feed (CDF) (CDF = CKD × DMI/d) and cost per kilogram of obtained gain (CKG) (CKG = CDF/ADG).

The data were subjected to homoscedasticity and normality tests, and analysis of variance was performed on all continuous variables with normal distributions. The initial weight was used as the covariant and, when not significant, was taken from the model. The mathematical model used was represented by:

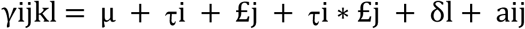

where *γ*ijkl = dependent variable; μ = overall mean; *τ*i = effect of factor i (inclusion level of the soybean hull); £j = effect of factor j (physical form of the corn); *τ*i*£j = interaction between factor i and factor j; δl = effect of initial weight; and aij = residual experimental error associated with the factorial inclusion level of the soybean hull and the physical form of the corn. The data were analyzed using t-tests at 5% significance to compare the means when the interaction of the studied factors was not significant (>5% significance).

## Results

No interaction was observed between corn processing and SH inclusion for the variables related to diet digestibility (Table 3). However, both the use of WC and the inclusion of SH reduced the TDN value of the diets (P = 0.02 and P = 0.03, respectively), although including SH increased the NDF content digested by the animals (P < 0.01). However, the DMAD, CPAD and NFCD results were not influenced by using WC or including SH, presenting averages of 0.87, 0.89 and 0.90, respectively.

**Table 3.**
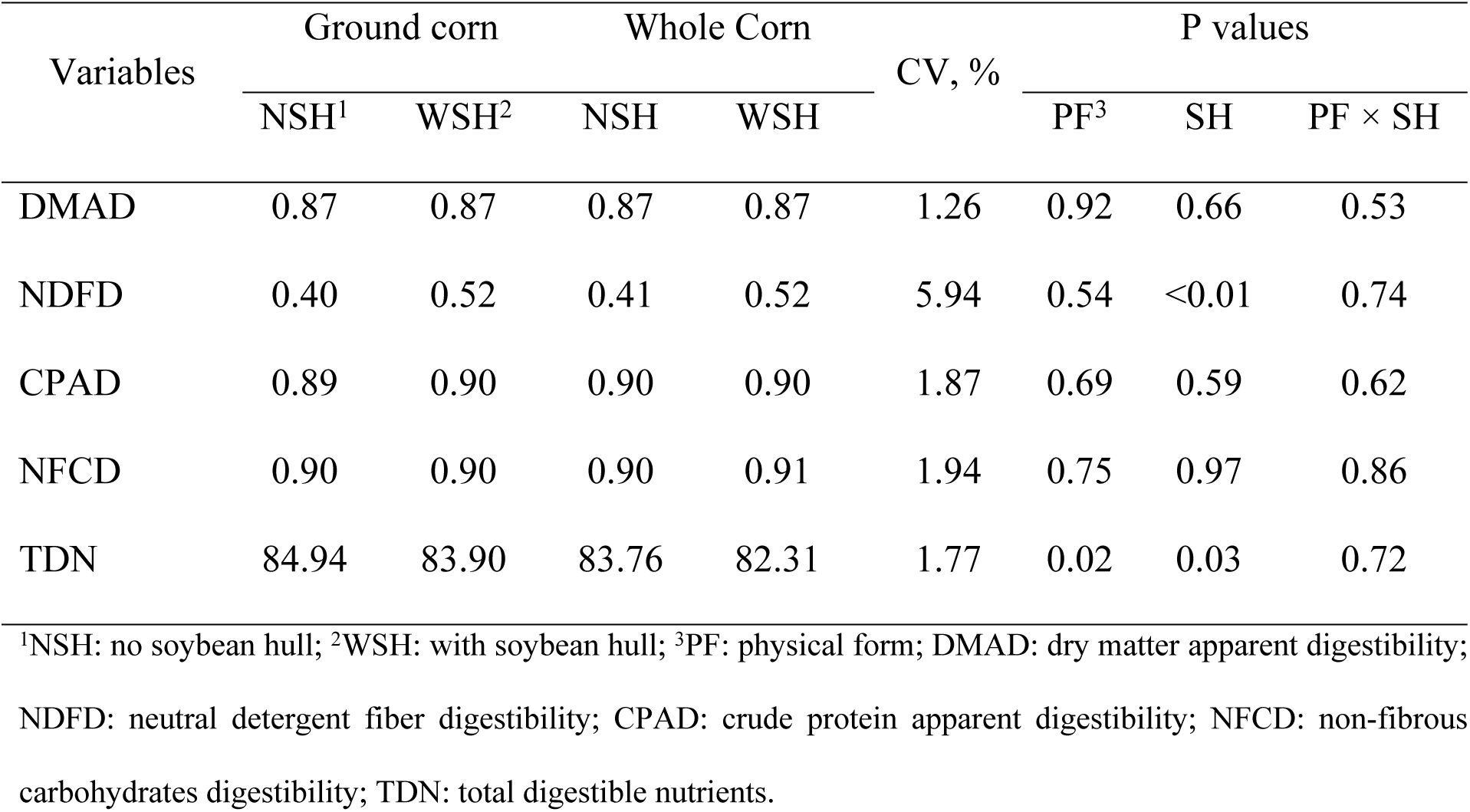
Apparent digestibility coefficients and total digestible nutrient values of the diets

No interaction was observed between the inclusion level of SH in the diet and the physical form of corn grain for the variables related to consumption (Table 4). However, when analyzed separately, SH use increased the NDFI (P < 0.01) and reduced the NFCI (P < 0.01), which was also reduced by using WC in the diets (P = 0.04). The other consumptions presented similar results, regardless of physical form of the corn grain or SH inclusion level, with means of 0.39 kg/d DMI, 0.88 kg/d TDMI, 0.07 kg/d CPI, 0.20 kg/d TCPI and 0.75 kg/d TDNI.

**Table 4.** Nutrient intake of crossbred dairy calves

| Variables,<br>kg/d | Ground corn |  | Whole Corn |  | CV, % | P values |  |  |
| --- | --- | --- | --- | --- | --- | --- | --- | --- |
|  | NSH<sup>1</sup> | WSH<sup>2</sup> | NSH | WSH |  | PF<sup>3</sup> | SH | PF × SH |
| DMI | 0.47 | 0.43 | 0.28 | 0.39 | 45.83 | 0.11 | 0.58 | 0.24 |
| TDMI | 0.96 | 0.92 | 0.77 | 0.89 | 20.22 | 0.12 | 0.56 | 0.25 |
| CPI | 0.09 | 0.07 | 0.06 | 0.08 | 49.06 | 0.38 | 0.87 | 0.23 |
| TCPI | 0.21 | 0.20 | 0.18 | 0.20 | 18.39 | 0.37 | 0.86 | 0.24 |
| NDFI | 0.05 | 0.21 | 0.02 | 0.20 | 55.93 | 0.50 | <0.01 | 0.76 |
| NFCI | 0.27 | 0.09 | 0.16 | 0.08 | 50.01 | 0.04 | <0.01 | 0.07 |
| TDNI | 0.82 | 0.78 | 0.65 | 0.73 | 21.4 | 0.08 | 0.73 | 0.29 |

<sup>1</sup>NSH: no soybean hull; <sup>2</sup>WSH: with soybean hull; <sup>3</sup>PF: physical form; DMI: dry matter intake; TDMI: total dry matter intake; CPI: crude protein intake; TCPI: total crude protein intake; NDFI: neutral detergent fiber intake; NFCI: non-fibrous carbohydrates intake; TDNI: total digestible nutrients intake.

The blood biochemical indicator results (Table 5) were not dependent on the evaluated factors, and the means were 88.57 mg/dl GL, 26.77 g/dl TCL, 36.82 mg/dl TG, 6.07 g/dl TP, 2.65 g/dl ALB, 6.33 mg/dl UR, 48.24 U/l AST, 116.19 U/l ALP and 1.19 mg/dl CRT.

**Table 5.** Biochemical indicators in the blood of crossbred dairy calves

| Variables | Ground Corn |  | Whole Corn |  | CV, % | P values |  |  |
| --- | --- | --- | --- | --- | --- | --- | --- | --- |
|  | NSH<sup>1</sup> | WSH<sup>2</sup> | NSH | WSH |  | PF<sup>3</sup> | SH | PF × SH |
| GL, mg/dl | 83.71 | 90.29 | 89.64 | 90.67 | 15.85 | 0.55 | 0.48 | 0.60 |
| TCL, mg/dl | 25.79 | 28.36 | 24.79 | 28.17 | 55.84 | 0.92 | 0.60 | 0.94 |
| TG, mg/dl | 28.93 | 40.00 | 42.36 | 36.00 | 33.43 | 0.32 | 0.62 | 0.07 |
| TP, g/dl | 5.70 | 6.13 | 6.49 | 5.98 | 11.57 | 0.23 | 0.88 | 0.09 |
| ALB, g/dl | 2.71 | 2.80 | 2.46 | 2.64 | 12.04 | 0.10 | 0.27 | 0.73 |
| UR, mg/dl | 6.57 | 5.86 | 8.07 | 4.83 | 42.07 | 0.81 | 0.06 | 0.22 |
| AST, U/ml | 46.03 | 47.14 | 44.17 | 46.13 | 24.18 | 0.42 | 0.99 | 0.69 |
| ALP, U/ml | 148.00 | 108.57 | 104.29 | 103.92 | 59.72 | 0.37 | 0.45 | 0.46 |
| CRT, mg/dl | 1.25 | 1.04 | 1.26 | 1.21 | 26.9 | 0.46 | 0.28 | 0.50 |

<sup>1</sup>NSH: no soybean hull; <sup>2</sup>WSH: with soybean hull; <sup>3</sup>PF: physical form; GL: glucose; TCL: total cholesterol; TG: triglycerides; TP: total protein; ALB: albumin; UR: urea; AST: aspartate aminotransferase; ALP: alkaline phosphatase; CRT: creatinine.

The physical form and the SH inclusion level no affected the variables related to the animals’ performance (Table 6). The means were 64.28 kg FW, 31.28 kg TWG, 0.56 kg/d ADG, 1.64 kg gain/kg MS in DME, 0.62 kg gain/kg MS in TDME, 8.69 kg gain/kg CP in CPE, 2.76 kg gain/kg CP in the TCPE and 0.75 kg gain/kg TDN in TDNE.

**Table 6.** Performance of crossbred dairy calves

| Variables | Ground corn |  | Whole Corn |  | CV, % | P values |  |  |
| --- | --- | --- | --- | --- | --- | --- | --- | --- |
|  | NSH<sup>1</sup> | WSH<sup>2</sup> | NSH | WSH |  | PF<sup>3</sup> | SH | PF × SH |
| IW | 32.93 | 32.00 | 32.43 | 34.67 | 18.75 | 0.65 | 0.78 | 0.51 |
| FW | 69.43 | 63.64 | 58.00 | 66.08 | 18.79 | 0.33 | 0.80 | 0.14 |
| TWG | 36.50 | 31.64 | 25.57 | 31.42 | 27.44 | 0.10 | 0.88 | 0.11 |
| ADG | 0.65 | 0.57 | 0.46 | 0.56 | 27.54 | 0.09 | 0.87 | 0.10 |
| DME | 1.51 | 1.52 | 2.00 | 1.54 | 38.11 | 0.29 | 0.35 | 0.32 |
| TDME | 0.68 | 0.61 | 0.58 | 0.63 | 19.92 | 0.48 | 0.85 | 0.20 |
| CPE | 8.00 | 9.79 | 9.27 | 7.73 | 43.60 | 0.79 | 0.93 | 0.26 |
| TCPE | 3.03 | 2.85 | 2.42 | 2.75 | 22.96 | 0.15 | 0.76 | 0.30 |
| TDNE | 0.80 | 0.72 | 0.69 | 0.77 | 20.00 | 0.62 | 0.99 | 0.18 |

<sup>1</sup>NSH: no soybean hull; <sup>2</sup>WSH: with soybean hull; <sup>3</sup>PF: physical form; IW: initial live weight in kg; PF: final live weight in kg; TWG: total weight gain in kg; ADG: average daily gain in kg/d; DME: dry matter conversion efficiency in kg/kg; TDME: total dry matter conversion efficiency in kg/kg; CPE: crude protein conversion efficiency in kg/kg; TCPE: total crude protein conversion efficiency in kg/kg; TDNE: total digestible nutrient conversion efficiency in kg/kg.

The data obtained in the morphometric evaluation presented results independent of the physical form of the corn and the CS inclusion level (Table 7) in evaluating both the final results and the respective gains during the suckling period.

**Table 7.** Morphometric measurements of crossbred dairy calves

| Variables,<br>cm | Ground corn |  | Whole Corn |  | CV, % | P values |  |  |
| --- | --- | --- | --- | --- | --- | --- | --- | --- |
|  | NSH<sup>1</sup> | WSH<sup>2</sup> | NSH | WSH |  | PF<sup>3</sup> | SH | PF × SH |
| IWH | 72.86 | 70.21 | 70.36 | 72.25 | 5.02 | 0.86 | 0.78 | 0.11 |
| IRH | 76.57 | 75.29 | 74.14 | 76.67 | 5.68 | 0.75 | 0.71 | 0.25 |
| IAP | 11.36 | 11.33 | 10.93 | 11.17 | 8.08 | 0.39 | 0.76 | 0.69 |
| IBL | 61.79 | 59.00 | 60.93 | 62.22 | 9.25 | 0.58 | 0.73 | 0.35 |
| ITP | 75.86 | 75.07 | 71.21 | 77.00 | 7.59 | 0.53 | 0.25 | 0.14 |
| WHG | 10.36 | 12.28 | 11.07 | 9.50 | 33.42 | 0.49 | 0.84 | 0.23 |
| RHG | 11.07 | 11.86 | 10.78 | 11.66 | 31.19 | 0.84 | 0.54 | 0.99 |
| APG | 1.00 | 1.24 | 0.93 | 1.00 | 46.78 | 0.41 | 0.38 | 0.70 |
| BLG | 12.36 | 11.71 | 10.65 | 8.95 | 47.50 | 0.29 | 0.60 | 0.74 |
| TPG | 20.07 | 19.86 | 15.28 | 18.25 | 39.39 | 24.71 | 0.61 | 0.60 |
| FWH | 83.21 | 82.50 | 81.43 | 81.75 | 5.45 | 0.46 | 0.91 | 0.76 |
| FRH | 87.64 | 87.14 | 82.36 | 88.33 | 7.11 | 0.38 | 0.25 | 0.18 |
| FAP | 12.36 | 12.57 | 11.86 | 12.17 | 6.83 | 0.16 | 0.42 | 0.88 |
| FBL | 74.14 | 70.71 | 67.43 | 71.17 | 7.37 | 0.13 | 0.94 | 0.08 |
| FTP | 95.93 | 93.50 | 86.50 | 95.25 | 9.01 | 0.71 | 0.56 | 0.73 |

<sup>1</sup>NSH: no soybean hull; <sup>2</sup>WSH: with soybean hull; <sup>3</sup>PF: physical form; IWH: initial wither height; IRH: initial rump height; IAP: initial arm perimeter; IBL: initial body length; ITP: initial thoracic perimeter; WHG: withers height gain; RHG: rump height gain; APG: arm perimeter gain; BLG: body length gain; TPG: thoracic perimeter gain; FWH: final wither height; FRH: final rump height; FAP: final arm perimeter; FBL: final body length; FTP: final thoracic perimeter.

Considering the cost of the ingredients during the evaluation period, SH and WC usages were insufficient for reducing feed costs (Table 8); the CDF was US $2.06, while the CQK was US $3.74.

**Table 8.**
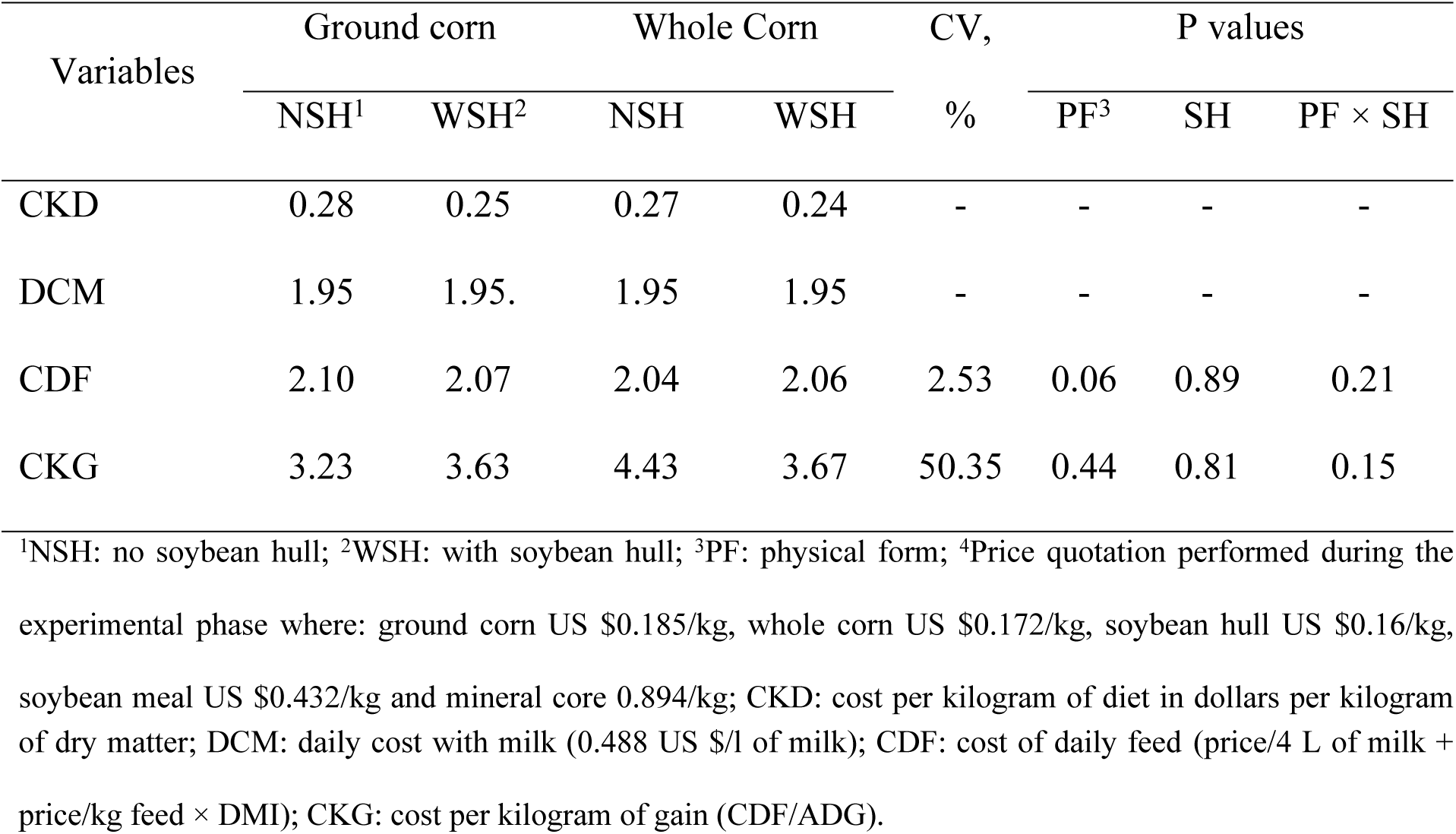
Costs associated with crossbred dairy calf feed

## Discussion

Supplying solid feed to calves beneficially affected prestomach development [4], as shown by the digestibility presented by the diets. The high digestibility observed for the dry matter and nutrients evaluated indicated that although the animals were young, using solid foods resulted in high digestibility of the nutrients in the diet, for example, CPAD, which is a nutrient highly required by animals during the initial production phase [16].

Although cattle digest fibrous feed efficiently only from the moment the rumen is functional, the high NDFD in WSH diets demonstrated that although this food is high in fiber (Table 1), the fiber has high quality because of the high pectin content [16] and low lignin content [16, 17]. This allowed even the young animals to take advantage of the fiber present in this feed, although at this age, they have a reduced capacity for digesting carbohydrates such as cellulose [18, 19], which is present in large quantities in SH (Table 1).

Although NDFD was favored, SH had a lower energy content than corn (3.4 Mcal/kg and 3.9 Mcal/kg, respectively [16]), causing it to reduce the TDN value of the diet. This result is common when CS is used instead of energetic concentrates, such as corn, in bovine diets [20, 21], and the literature shows that as the SH inclusion level increases, expressive variations in the TDN value of the diet increases [9].

This reduction in the amount of digestible nutrients available to the animals also occurred with WC grain use, despite the diets with whole and ground grain presenting the same chemical-bromatological compositions (Table 2). However, this variation between the TDN level in the diets with WC or GC was only 1.64%, representing minimal change. Thus, the proximity between the TDN value of the diets with WC and GC allowed the animals fed these diets to revert to the difference in TDN value, since they achieved TDNI results similar to those fed GC.

This similarity was evidenced in the animals’ blood data, since the glycemic levels remained within the range considered normal for the animal category [22], indicating that the animals received sufficient nutrients to meet their needs. In addition, the creatinine, AST and ALP results indicated that the experimental diets did not alter or impair the animals’ hepatic (AST and ALP) [23,24] or renal (creatinine) functions [25,26]. That is, the diets promoted little metabolic challenge to the suckling calves, as the levels of these indicators remained within the normal variation ranges for suckling Holstein calves per [27] (creatinine between 0.98 and 1.56 mg/dl, AST between 27.6 and 46.3 U/ml and APL between 67.71 and 141.33 U/ml).

Considering the intake of other nutrients, the corn grain used in this work was the hard type (flint) associated with the use of nonpelleted feed, such as soybean meal, and the mineral core may have contributed to the reduced NFCI, since the animals fed whole grains selected their diets, preferentially feeding on components of the diet that were easier to chew, to the detriment to corn grain consumption. This selection favored the decreased NFCI since the soybean meal had a much lower NFC content than that found in corn (184 and 754 g/kg, respectively [16]).

In addition, because soybean hulls have NFC contents lower than that found in corn, as well as high NDF levels (Table 1) and are constituted basically by structural carbohydrates, their inclusion in the diet also contributed to the reduced NFCI, resulting in an increased NDFI. Results similar to those in the present study are commonly reported in the literature when SH is included in diets that present high energy contents [28, 29]. This occurs because SH inclusion usually reduces the proportion of foods, such as corn and sorghum, which present high NFC contents and low NDF ratios.

DMI and TDMI (concentrate + milk) occurring similarly among the diets indicated that the diets were well-accepted by the animals and showed good palatability. These characteristics are very important at this stage of production, since ingesting solid food reduces the animals’ dependence on liquid food [30], enabling the weaning age to be reduced [16] and accelerating the return of invested capital in animal production.

In addition, the DMI and TDMI variables were 0.39 and 0.89 kg/d, respectively. These results were lower than those observed in the work of [24], which evaluated the inclusion of soybean hulls in the diet of calves during the breeding phase and found a mean DMI of concentrated feed of 0.49 kg/d. However, despite the lower consumption values, the feed conversion results were better, and the animals needed to consume 1.70 kg of TDM to gain one kilogram live weight, whereas in the work by [31], the animals needed 1.91 kg/d, showing that the animals were highly efficient, what corresponded to a reflex in the weight gain, since the animals presented ADG of 0.56 kg/d.

We compared the results of weight gained in the WSH diets with the results of other studies in the literature, and found that SH has a superior performance during the rearing phase (ADG of 0.35 [32] and 0.45 kg/d [33]) when evaluating by-products similar to SH with high fiber content and low particle size, such as babassu mesocarp meal [32] and citrus pulp [33], in the diets of dairy calves. Thus, SH is a good option to be used in facilities that produce this type of animal, since it allows the animals to be weaned with high weights.

The animals fed the experimental diets developed well based on the data from the morphometric measurements performed on the animals during the suckling phase. During the evaluation period, the animals presented significant gains for these measurements; however, these gains were similar between the diets, which reflected the similarity in the animals’ weight gain and TDNI. This was also highlighted by another study [34] that obtained similar metabolizable energy intakes in Holstein calves, thus resulting in similar morphometric measurements for these animals.

Despite the favorable results regarding animal performance, SH use or the reduction in processing by using WC did not reduce feed costs, even when the costs were analyzed as a function of weight gain. These results demonstrate the need for a greater difference in the market value between GC and WC and between GC and SH to significantly reduce production costs by using WC or SH in calf diets during suckling.

Reduced diet costs with SH use was observed by [35], who observed a 20% decrease in costs per kilogram weight gain by using up to 30% SH in Nellore steer diets. However, in this study, the difference between the GC and SH prices was 25%, whereas in the present study, the variation was only 14%.

In addition, when considering calf production costs, milk represents a large proportion of daily feed costs (94%), demonstrating the importance of reducing the animals’ weaning age. As milk is the main source of income for dairy farms, reducing the weaning age reduces calf production costs, as it reduces the amount of milk used to feed the animals during rearing [36].

## Conclusion

Using soybean hull or whole corn in crossbred dairy calves’ diets during the rearing phase does not impair their performance or change their blood markers; however, it does not reduce the costs associated with feed during rearing.

## Acknowledgments

The financial support provided by CNPq for the realization of this research.

